# Improved genome assembly of whale shark, the world’s biggest fish: revealing “chromocline” in intragenomic heterogeneity

**DOI:** 10.1101/2025.09.06.674125

**Authors:** Yawako W. Kawaguchi, Rui Matsumoto, Shigehiro Kuraku

**Affiliations:** Molecular Life History Laboratory, National Institute of Genetics, Mishima, Shizuoka, 411-8540, Japan; Okinawa Churashima Research Center, Okinawa Churashima Foundation, Okinawa, Japan; Okinawa Churaumi Aquarium, Okinawa, Japan; Department of Genetics, Sokendai (Graduate University for Advanced Studies), Mishima, Shizuoka, Japan; Laboratory for Phyloinformatics, RIKEN Center for Biosystems Dynamics Research, Kobe, Hyogo, 657-0024, Japan

## Abstract

High-quality chromosome-level assemblies are essential for understanding genome evolution but remain difficult to obtain for large and complex genomes. Here we present a near gap-free genome assembly of the whale shark (*Rhincodon typus*) generated with long-read sequencing and Hi-C scaffolding, markedly improving contiguity and completeness. In particular, the X chromosome was extended to nearly twice its previous length, and putative pseudoautosomal regions were identified. Moreover, we report the first Y-linked scaffolds for this species. Comparative analyses with the zebra shark revealed exceptionally low substitution rates across the genome. We further detected a negative correlation between chromosome length and synonymous substitution rate (*d*_*S*_), explained by a positional gradient, designated as “chromocline”, in which substitution rates gradually decrease from chromosomal ends toward central regions. Notably, the X chromosome exhibited low *d*_*S*_ compared with autosomes of similar size, consistent with male-driven evolution. Our results highlight positional and sex-chromosome effects as key determinants of molecular evolutionary rates. The improved assembly will enable broad application to population-genetic and conservation genomic analyses in the whale shark.

## Background

Achieving complete and accurate genome assemblies remains a significant challenge in genome informatics. It is hindered primarily by intrinsic genomic complexities such as large size, high repetitiveness, and heterozygosity, compounded by limitations in sequencing data quality and quantity. Technological advancements and accumulated experience are making complete genome sequencing more accessible [1], but in reality, genome assembly is often an iterative process, leading to multiple versions for a single species, frequently generated by diverse research entities. Existing guidelines largely focus on initial assembly finalization [2], often neglecting critical considerations for releasing improved versions to avoid traceability issue, particularly regarding consistency of chromosome and sequence identifiers. This oversight severely impedes data interoperability, reusability, and reproducibility. Thus, recommended practices for iterative genome assembly improvement and the subsequent formal release of updated versions remain to be formulated.

The species chosen in this study, whale shark *Rhincodon typus*, is the largest ‘fish’ species among extant fish lineages encompassing jawless, chondrichthyan, and osteichthyan species. The whale shark is categorized as endangered (EN) in the IUCN Red List, which limits tissue sampling for biological studies. Genome sequencing for this species was initiated with assembling short reads by Read et al. to obtain highly fragmented contigs [3]. This study used a postmortem male tissue sampled at Georgia Aquarium. Later, Hara et al. performed reassembly of the short reads obtained by Read et al [3,4], and scaffolded the contigs with mate pair reads produced using blood cell DNA with mate-pair library preparation protocol iMate [5]. In parallel, Weber et al. obtained a heart tissue from a deceased male at Hanwha Aquarium, Jeju, Korea, and combined short reads and mate-pair, with TruSeq Synthetic Long Read (TSLR) libraries to obtain the assembly RhiTyp_1.0 [6]. In 2021, the residual tissue sample used by Read et al. was used for obtaining consensus long reads (CLR) with single molecule, real-time (SMRT) technology of Pacific Biosciences [3]. The first release of chromosome-scale assemblies was achieved by DNA Zoo consortium which scaffolded the above-mentioned assembly sequences by Weber et al. with Hi-C data prepared with male tissues (https://www.dnazoo.org/assemblies/rhincodon_typus). Most recently, Yamaguchi et al. utilized the 10X Chromium Linked Read data [7] as part of data production in the Squalomix consortium [8]. The contigs resulted from this data were scaffolded by blood cell Hi-C data prepared with the iconHi-C protocol [9]. This assembly sRhiTyp1.1, is derived from the sexually mature male individual (named Jinta) maintained at Okinawa Churaumi Aquarium since 1995 [10] and is labelled as ‘reference’ at NCBI Genomes, as of July 2025.

Recent advances have enabled the identification of sex chromosome sequences in multiple cartilaginous fishes, a group known to exhibit male heterogamety [11]. While contiguous X chromosomes have been reported for some species, their completeness and taxonomic coverage remain limited [7,12,13]. Y chromosomes, often highly repetitive and small, are even more elusive; only a few reports have constructed partial sequences identified to date [12,13]. Notably, sex chromosomes of sharks and rays are thought to have been conserved for over 300 million years, underscoring the evolutionary importance of expanding genomic insights into these chromosomes [13]. In whale shark, an X chromosome was previously identified by Yamaguchi et al. (2023), but this effort resulted in relatively short (∼12 Mbp) and fragmentary scaffold, compared with the ∼20 Mbp-long X chromosome of its close relative, the zebra shark. No sequences from the Y chromosome have been reported for this species [7]. Thus, more complete X chromosome assemblies and the discovery of Y-linked sequences across broader taxa are essential to understand sex chromosome evolution in this lineage.

Resolving chromosome-scale assemblies including the sex chromosomes provides a window into how genomic position shapes evolutionary rates. Gene evolutionary rates are not solely determined by gene function but are also influenced by their genomic location [14–16]. For example, in the sex chromosome context, male-driven evolution predicts lower neutral substitution on the X chromosome than on autosomes [17–19], whereas hemizygosity can accelerate adaptive change on X chromosomes (faster-X effect, [20]). Beyond sex linkage, both birds and mammals have shown faster evolution on shorter chromosomes [21–23]. One theoretically grounded explanation is that the obligatory crossover elevates recombination per unit length on small chromosomes, and recombination-associated break/repair together with linked selection or GC-biased gene conversion can raise local substitution rates [15,24]. Positional effects also occur within chromosomes; in Xenopus, synonymous divergence increases with distance from centromeres and tends to be higher toward chromosomal ends [25]. These patterns collectively motivate joint tests of genomic location, chromosome size, and sex linkage. However, strong intra-chromosomal heterogeneity in recombination further complicates inference, and comprehensive analyses that disentangle these factors remain scarce outside Tetrapoda.

In this study, we present a new chromosome-level genome assembly for the whale shark. Our assembly reveals a more complete X chromosome and, for the first time, putative Y chromosome sequences. Leveraging this new resource, we test the influence of chromosome length, chromosomal position, and sex linkage on synonymous substitution rates. These analyses reveal positional and sex-chromosome effects on molecular evolutionary rates, providing a foundation for future comparative studies.

## Results

### Long-read-based chromosome-scale genome assembly of whale shark

We generated a new genome assembly for the whale shark (*Rhincodon typus*) by conducting long-read sequencing of an adult male individual, followed by *de novo* assembly with Hi-C scaffolding (Figure 1A, B). Our new assembly (named sRhiTyp1.2) has a total size of 3.22 Gbp and consists of 3,201 scaffolds. Compared to two previous versions (RhiTyp_1.0 and sRhiTyp1.1), which were 2.82 Gbp in size with 136,451 scaffolds and 2.88 Gbp in size with 16,776 scaffolds, respectively, the new assembly exhibits a marked improvement in contiguity (Figure 1C, Table S1) [6,7] Furthermore, the assembly size of sRhiTyp1.2 now approximates the estimated genome size of 3.75 Gbp previously measured with flow cytometry [1]. Notably, compared to sRhiTyp1.1, the number of undetermined bases decreased remarkably (95,223 to 470), and the BUSCO completeness score strikingly increased (84.2% to 97.9%, Figure 1B, Table S1). These enhancements indicate a more complete and chromosome-level assembly.

**Figure 1.**
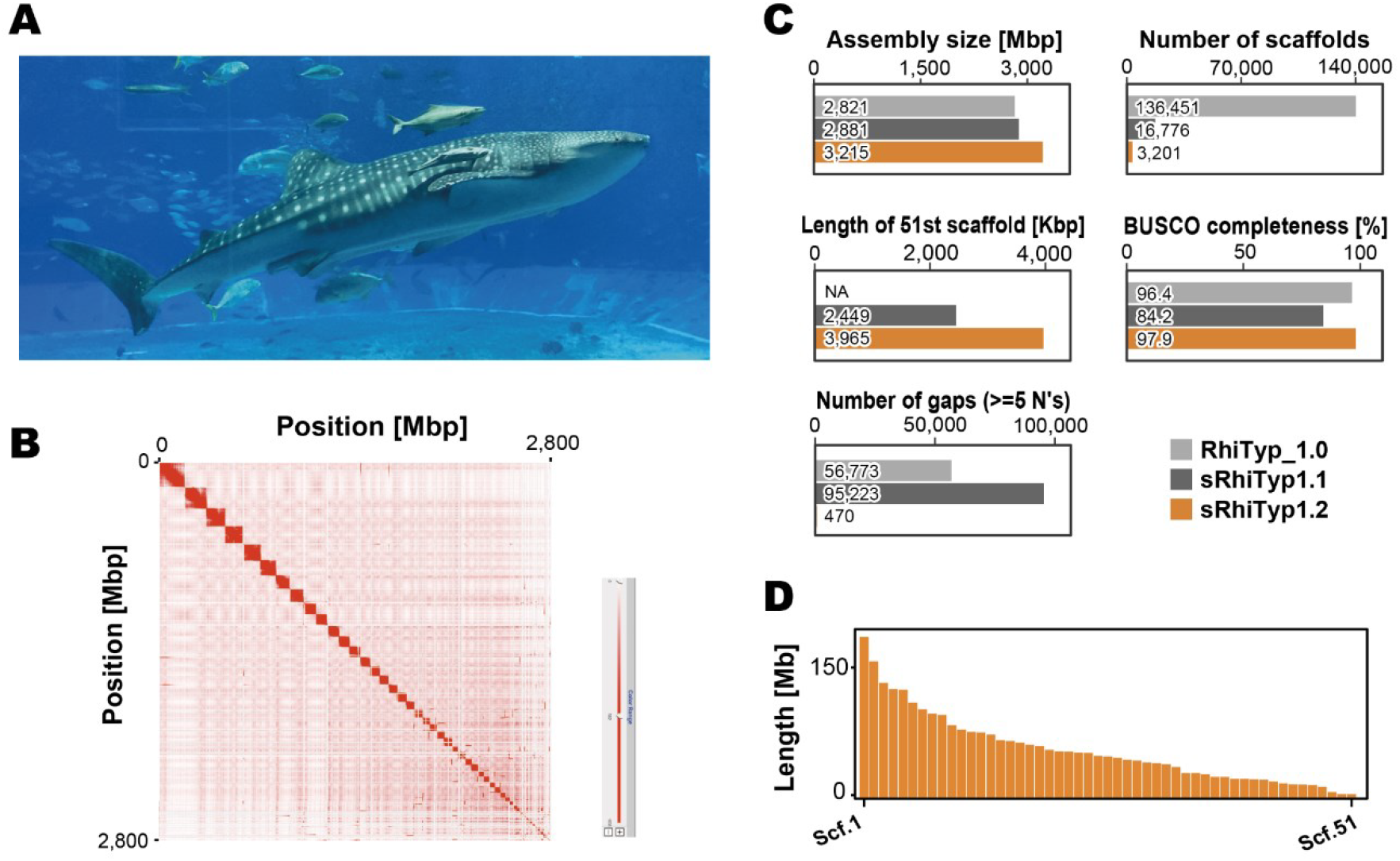
Statistics of the new genome assembly of whale shark (Rhincodon typus). (A) Whale shark individual, named Jinta, from which a genome assembly was obtained in this study. Photo credit: Akifumi Yao. (B) Hi-C contact map of top 51 scaffolds visualized using Juicebox, showing chromatin contact profiles supporting chromosome-scale scaffolding. (C) Comparative assembly statistics between the previous versions (RhiTyp_1.0 and sRhiTyp1.1) and our new version (sRhiTyp1.2). (D) Length distribution of the 51 longest scaffolds, defined as the chromosomes. Abbreviations: Scf, scaffold; Chr, chromosome.

We defined the putative chromosome set by selecting the 51 largest scaffolds (Figure 1D), based on the known chromosome number of the whale shark [11], and designated them as chromosome-scale sequences for downstream analyses. Within this set, the smallest chromosome—scaffold 51—was extended from 2.45 Mbp to 3.97 Mbp compared to sRhiTyp1.1. The chromosome lengths vary from 3.97 Mbp to 185 Mbp, which reflects a typical length spectrum observed in cartilaginous fishes [7,26].

### More complete sex chromosome sequencing

To identify sex chromosomes, we mapped short-read whole genome sequencing data from both male and female individuals to the new assembly. Among the 51 scaffolds presumed to represent chromosomes, chromosome 40 exhibited a male-to-female read depth ratio of approximately 1:2, a hallmark of the X chromosome in a species of male heterogamety. Based on this pattern, we designated it as the X chromosome (Figure 2A). The X chromosome in the new assembly was nearly twice as long as in the previous version (increased from 12 Mb to 21 Mb; Fig 1B, 2B), and the number of annotated genes increased from 247 to 281. Regions at both ends of the X chromosome exhibited similar depth between sexes (Figure 2C), suggesting the presence of pseudoautosomal regions (PARs), in which recombination may occur between X and Y chromosomes.

**Figure 2.**
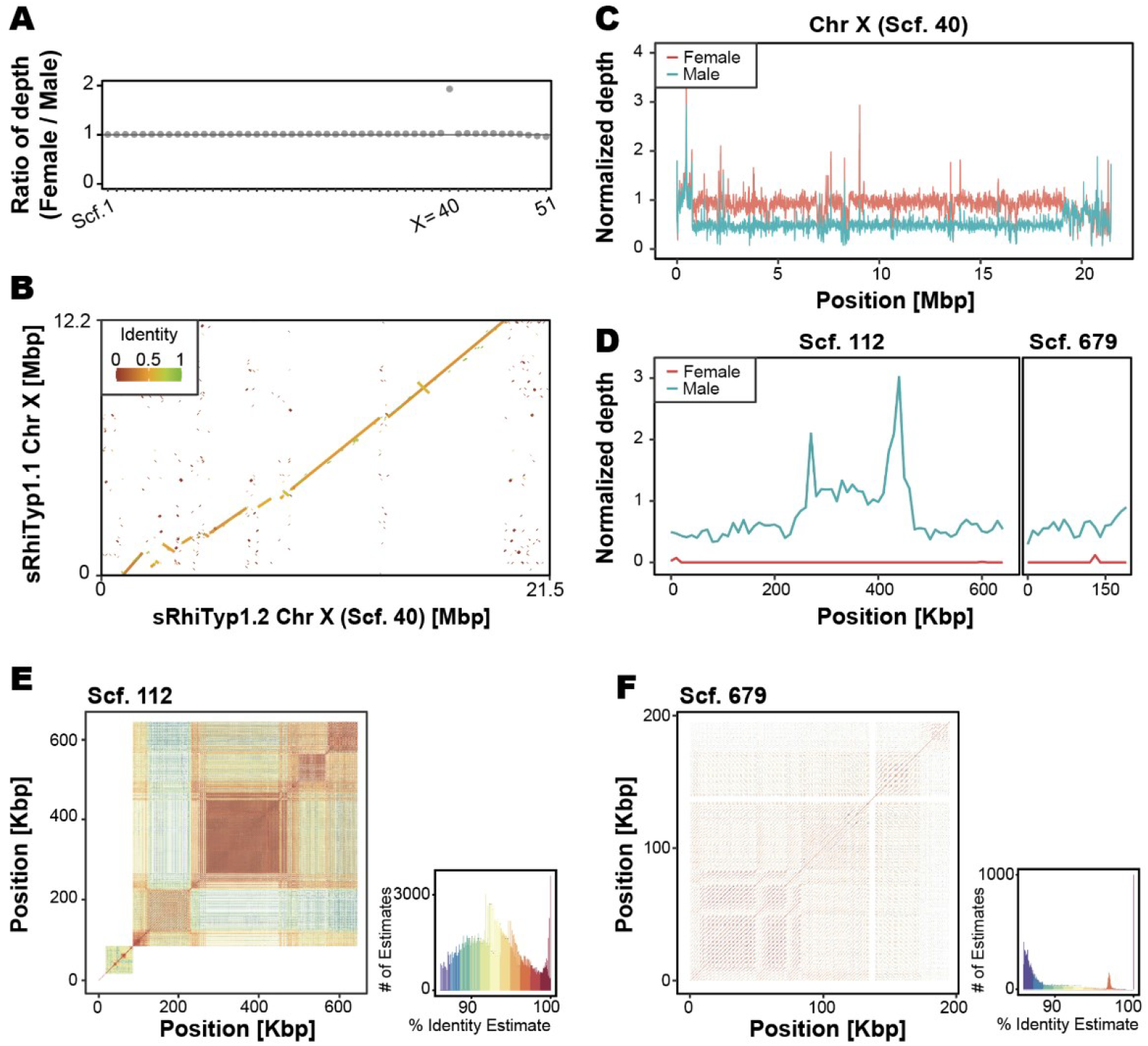
Identification and characterization of sex chromosomes. (A) Female to male ratio and median of the depth of genome short reads for each chromosome. Scaffold 40 displays approximately twice the read depth in females compared to males, consistent with an X chromosome. (B) Dot plot of the X chromosome comparing the new assembly with the previous assembly (sRhiTyp1.1). (C) Depth distribution in the X chromosome. (D) Depth distribution in the two putative Y chromosome fragments. (E, F) Identity heatmaps for the two putative Y chromosome fragments. Abbreviations: Scf, scaffold.

We further searched for Y-linked scaffolds by identifying regions with high read depth in males but low or no coverage in females. This approach led to the identification of two scaffolds with the differential mapping results between the sexes that are presumed to be fragments of the Y chromosome: scaffold0112 (645,398 bp) and scaffold0679 (195,000 bp) (Figure 2D). Each of these two scaffolds contained only a few predicted genes (two and one, respectively), and none of those genes were found to be implicated in sex determination documented in other species so far. Because scaffolds with male-restricted coverage could in principle result from contaminants specific to the sequenced male individual, we verified their origin by BLAST searches against the NCBI nr database. The predicted genes exhibited strong matches to homologs from other elasmobranch species, demonstrating that these scaffolds represent genuine shark sequences rather than contaminants. Their sequence composition was characterized by a high abundance of repeats (Figure 2E, F). This repeat-rich and gene-poor pattern is consistent with previous observations of Y chromosomes in other shark species [13].

### Substitution rate variation within and between chromosomes

To assess regional patterns of molecular evolution, we quantified substitution rates by comparing orthologous protein-coding genes between the whale shark and the zebra shark (*Stegostoma tigrinum*), the closest extant relative of the former species. Because some chromosomes harbor only a few orthologs, which can yield unstable median estimates, we summarized per-chromosome rates only for chromosomes containing more than ten orthologs. Across these chromosomes, substitution rates varied appreciably: the median values of synonymous substitution (*d*_*S*_) ranged from 0.0612 to 0.111, non-synonymous substitution (*d*_*N*_) from 0.0153 to 0.0339, and *d*_*N*_/*d*_*S*_ from 0.173 to 0.326 (Fig. 3A). Notably, *d*_*S*_ showed a significant negative correlation with chromosome length, while *d*_*N*_ exhibited a weaker but still negative trend (Figure 3A), pointing to an effect of chromosome length on substitution rates. Accordingly, shorter chromosomes show higher neutral substitution rates, with only a modest increase in nonsynonymous rates.

**Figure 3.**
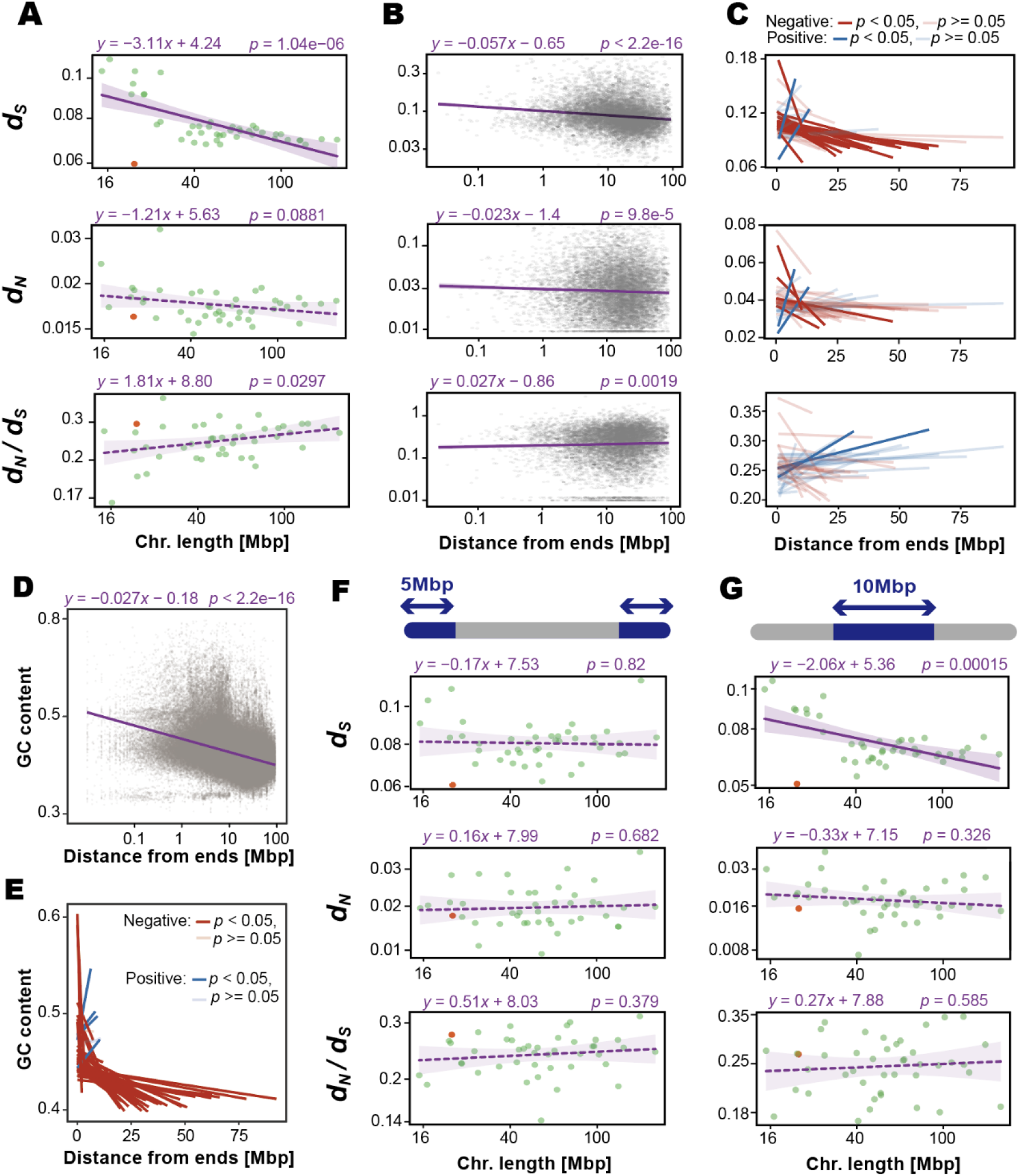
Substitution rates across chromosomal regions. (A) Chromosome lengths plotted against the median substitution rates for chromosomes containing more than 10 genes. (B) Substitution rates of individual genes plotted against their distance from chromosome ends. (C) Slope estimates from regressions between substitution rates and distance from chromosome ends, calculated separately for each chromosome. (D) GC content in 10-kbp windows plotted against their distance from chromosome ends. (E) Slope estimates from regressions between GC content and distance from chromosome ends, calculated for individual chromosomes. (F) Chromosome lengths plotted against the median substitution rates, for only genes located within 5 Mbp of chromosome ends. (G) Chromosome lengths plotted against the median substitution rates, for only genes located within 10 Mbp-long stretches at the centers of chromosomes. In A, F, and G, Green and orange dots indicate autosomes and the X chromosome, respectively. Solid and dashed lines indicate significant (*p* < 0.05) and non-significant simple regressions, respectively; shaded areas represent 95% confidence intervals. All axes in A, B, D, F, and G are shown on logarithmic scales.

To scrutinize this pattern, we analyzed the relationship between gene position (distance from chromosome ends) and substitution rate. We found that genes located closer to the ends tended to exhibit higher *d*_*S*_ and *d*_*N*_, significantly (Fig. 3B). At the individual chromosome level, 35 out of the 44 chromosomes showed a negative correlation between *d*_*S*_ and distance from the chromosome ends, 19 of which were statistically significant, including the X chromosome (Fig. 3C, *p* < 0.05). In contrast, only nine chromosomes showed a positive correlation, with only two of them showing significance. GC content also negatively correlated with distance from chromosome ends (Figs. 3D, E). At the individual chromosome level, most chromosomes also displayed negative correlations. Together, these patterns indicate pronounced intrachromosomal heterogeneity, manifesting as a positional cline with distance from chromosome ends.

The intrachromosomal heterogeneity could underlie the chromosome-length effect, because shorter chromosomes harbor a larger fraction of end-proximal, fast-evolving genes than larger chromosomes do. We therefore asked whether the length effect is independent of intrachromosomal heterogeneity, specifically whether it persists at fixed intrachromosomal position. Focusing on only genes within 5 Mb of the ends renders the length effect undetectable (Fig. 3F). It indicates that the length-dependent variation in nucleotide substitution rates is largely driven by gene positioning along the chromosome. In contrast, focusing on only genes located in the central regions of chromosomes (within the central 10 Mbp), the length effect persisted (Fig. 3G). This suggests that the variations in substitution rates are better explained by proximity to chromosome ends than to central regions—supporting a model in which distance from the ends rather than from the center is the primary positional driver of intrachromosomal substitution rate variation.

An exception to this overall pattern was observed in the X chromosome (scaffold 40). Although it is one of the relatively short chromosomes (40th out of 51), it exhibited significantly lower *d*_*S*_ compared to autosomes (Figs. 3A and 4A). *d*_*N*_ was not significantly different, and *d*_*N*_/*d*_*S*_ was slightly but not significantly elevated on the X chromosome. Notably, this reduction in *d*_*S*_ remained significant even when only telomere-proximal genes were analyzed (Figs. 3F and 4B), suggesting that the X chromosome has intrinsically lower substitution rates, independent of gene position. This pattern is consistent with a male-driven evolution in this species. To evaluate this more directly, we sought to estimate *d*_*S*_ for the Y chromosome; however, the putative Y scaffolds contained too few genes to support reliable rate estimates and were therefore excluded.

**Fig 4.**
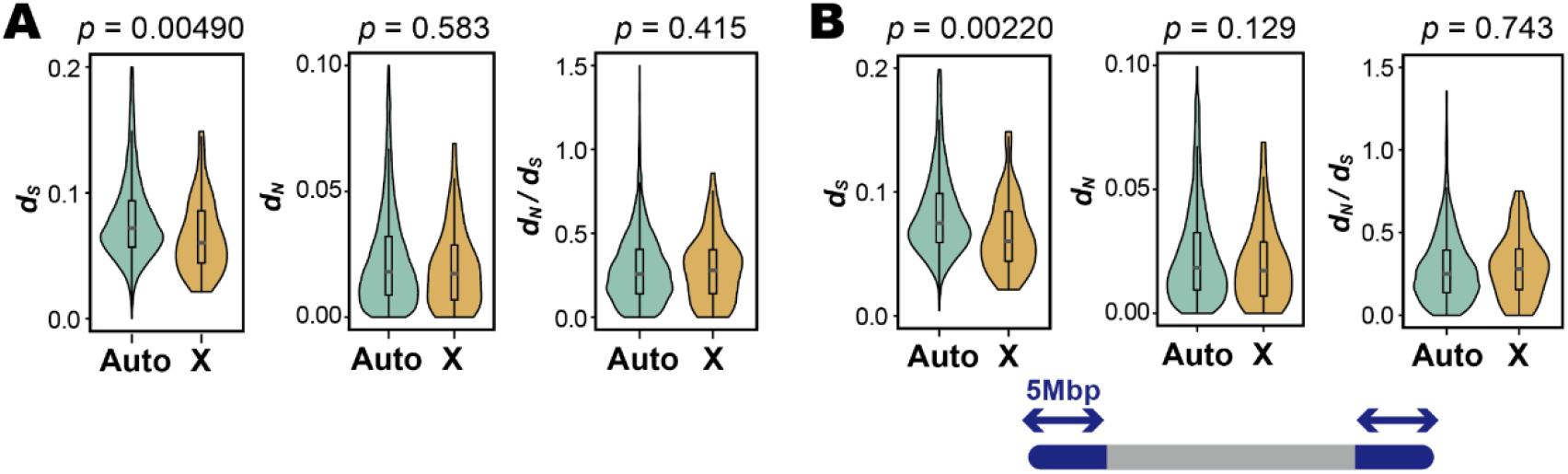
Substitution rates of each chromosome. (A) Comparison of substitution rates between autosomes and the X chromosome using all genes. (B) Focusing on the only genes within 5 Mbp of chromosome ends. Bars and boxes represent the median and interquartile range, respectively; whiskers extend to 1.5 times the interquartile range. Statistical significance was assessed using a randomization test with 10,000 permutations. Abbreviations: Auto, autosome; X, X chromosome.

## Discussion

This study provided a new genome assembly sRhiTyp1.2 of the world’s biggest fish, whale shark. It also serves as a case study for a frequent issue in modern genomics: the release of a new genome assembly for a species that already has one. Crucially, its importance is infrequently emphasized in scientific publications. Considering the coherence between different versions, we adjusted the nucleotide sequence orientations of the individual chromosomes to those in the previous assembly sRhiTyp1.1. Importantly, this is essential for confirming the location of genes and other genomic elements and avoiding traceability issues. In fact, maintaining consistent chromosome identifiers is dependent on successfully assembling and identifying complete chromosomes. Consideration on these two items is demanded in finalizing the assembly sequence set to be released in public. However, successful retrieval of individual chromosomes, even for relatively small ones, is a prerequisite of maintaining the chromosome identifiers. In our present study, scaffold sequences in the novel assembly sRhiTyp1.2 were sorted by their lengths and given new identifiers according to their lengths. This is because some of the scaffolds in the previous assembly version were supposedly fragments of large chromosomes but not entire chromosomes. When complete sequences become available, further reorientation should be considered based on centromere positions to designate the end of individual short chromosome arms as the beginning of the chromosomal sequences.

Our new whale shark assembly substantially reduces gaps and extends the X chromosome, yielding far more complete chromosome resolution than prior versions (Fig 1C and 2B). Although previous efforts had achieved near chromosome scale assemblies using short reads combined with Hi-C scaffolding, the contigs in those assemblies were relatively short, necessitating the insertion of large gaps during scaffolding [7]. In contrast, our approach utilized long-read sequencing with Oxford Nanopore technology, which dramatically reduced the number of gaps and improved assembly contiguity. This enabled more precise comparative analyses with the closely related zebra shark, whose genome was previously studied at the chromosome level. In addition, we identified, for the first time in this species, two putative scaffolds from the Y chromosome (Fig 2D, E and F). The gene repertoires on these scaffolds were extremely sparse, consistent with previous reports of the Y chromosomes of other shark species, and the high repeat content resembled that of the bamboo shark, a close relative [13]. Furthermore, the X chromosome in our assembly was nearly twice as long as the counterpart in the previous version, and we identified putative PAR at two ends, which were not detected in earlier assemblies.

The divergence between whale shark and zebra shark is estimated at approximately 50 million years ago [27]. We found that the median synonymous substitution rates (*d*_*S*_) across chromosomes ranged from 0.0612 to 0.111, suggesting exceptionally slow rates of molecular evolution (Fig 3A). For comparison, the human–mouse median *d*_*S*_ is approximately 0.58 over 90 million years [28], the chicken–zebra finch *d*_*S*_ is ∼0.4 over 66–86.5 million years [29,30], and the Tetraodon–Takifugu *d*_*S*_ is ∼0.59 over 32–55 million years [30,31]; reviewed in [32]. These comparisons underscore the exceptionally slow molecular clock of these shark lineages.

While we observed a negative correlation between chromosome length and synonymous substitution rate aligning with prior studies (Fig 3A) [21–23], we also demonstrate that the correlation is largely explained by length-independent intrachromosomal heterogeneity (Fig. 3B, C, F and G). We term this heterogeneity “chromocline”, in which sequence disposition changes gradually from chromosome end (Fig. 5). When a single representative rate per chromosome (e.g., median or mean across genes) is computed, it interacts with gene position. Longer chromosomes contain a greater fraction of genes at larger absolute distances from telomeres, which depresses their chromosome-level summaries. In contrast, shorter chromosomes are enriched for telomere-proximal genes, which elevates their summaries. This compositional effect produces the apparent negative correlation between chromosome length and synonymous substitution rate. Consistent with a shared positional mechanism, GC content exhibits a parallel cline, decreasing with distance from chromosome ends (Fig. 3D and E). This coupled gradient in substitution rates and base composition strongly supports a sequence architecture aligning “chromocline”. This result demonstrates that the length dependence of substitution rates is not an intrinsic property of chromosome size, but rather reflects the distribution of genes with respect to distance from the ends. Notably, a similar gradient of synonymous substitution rate has been reported in *Xenopus* [25], raising the possibility that the widely observed chromosome-length effect reflects a conserved “chromocline”.

**Fig. 5.**
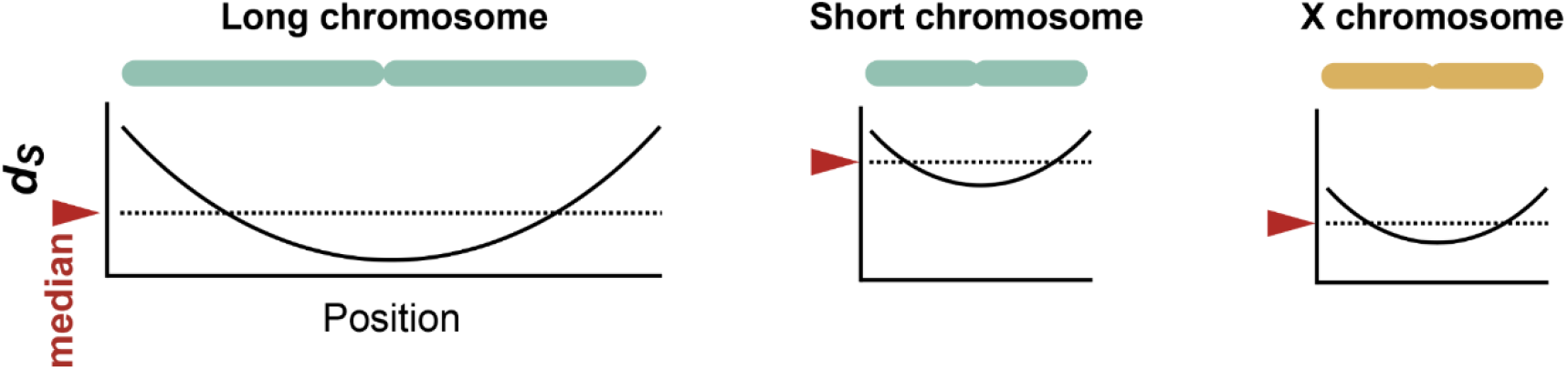
Schematic diagrams of the *d*_*S*_ distribution along chromosomes, “chromocline”. Longer chromosomes have more genes located far from the ends, resulting in a lower overall median *d*_*S*_ (red arrows). Conversely, shorter chromosomes tend to have higher median *d*_*S*_ because they have fewer genes located far from their ends. The X chromosome, however, exhibits unusually low median *d*_*S*_ despite its small length.

Such elevated rates near telomeres may reflect increased recombination activity in subtelomeric regions—a pattern commonly reported in animals [33,34]. Alternatively, or in addition, chromatin state, and DNA repair mechanisms may differ between distal and central chromosomal regions, contributing to the observed variation [35,36]. The consistent effect of this “chromocline” across many chromosomes in the whale shark genome provides an explanation for the inter-chromosomal variation in nucleotide substitution rates. A few chromosomes showed higher synonymous substitution rates toward their centers. This heterogeneity may reflect chromosome-specific landscapes in recombination or chromatin state. However, since these chromosomes were also among the shortest, we cannot exclude possibility that they represent assembly fragments of larger chromosomes, so we interpret these slopes cautiously.

Interestingly, despite being one of the relatively short chromosomes (40th out of 51), the X chromosome exhibited significantly lower synonymous substitution values than autosomes of similar length (Fig. 3A and 4). While we cannot entirely rule out residual uncertainty at the terminal regions of X, prior cytogenetic and genomic studies in sharks generally report short-to-intermediate X chromosomes, rendering a large error in X-chromosome length unlikely [11,12]. This pattern persisted even when we examined only telomere-proximal genes, indicating that the reduction in synonymous substitution rates on the X chromosome is not due to gene positioning but rather reflects its chromosome-wide characteristic (Fig. 3F and 4B). This is consistent with the male-driven evolution hypothesis: the mutation rate is higher in the male germline, and X chromosome spend proportionally less time in males, so neutral substitutions accumulate more slowly on the X chromosome [17–19]. Conversely, we found no clear evidence for Faster-X evolution, suggesting that adaptive evolution is not particularly accelerated at least in protein-coding genes. One potential explanation is the absence or incompleteness of dosage compensation in sharks. It has been proposed that the efficacy of Faster-X evolution may depend on the degree of dosage compensation [37,38] and sharks have recently been reported to lack complete dosage compensation [12,13], which could attenuate the signal of positive selection on the X chromosome. In case of birds (ZZ/ZW), dosage compensation is generally incomplete on the Z chromosome, and many avian clades show Faster-Z patterns often attributed to reduced effective population size with contributions from positive selection and male-biased mutation can elevate neutral rates on Z [39,40]. Taken together, male mutation bias, hemizygosity, and dosage-compensation regimes determine how sex-chromosome signals vary among lineages.

Our results clarify that chromosome-length effects on substitution rates largely reflect a genome-wide, position-dependent “chromocline”. We infer that recombination rate heterogeneity underlies much of the cline though direct recombination maps in whale shark will be required to test this. In parallel with this recombination rate view, interpreting the X-specific slowdown requires resolving its counterpart, the Y chromosome. As in other sharks, the Y chromosome of the whale shark appears gene-poor and repeat-rich, rendering sequence recovery challenging. However, in other words, the gene-poor Y chromosome, together with the overall slow substitution rate and a karyotype with remarkably chromosome length variations, is a distinctive hallmark of sharks. These features position sharks as a powerful model to understand how genomic position, in concert with sex linkage, shapes the tempo and mode of gene evolution.

## Materials and Methods

### Animal tissue sampling

Fresh blood from a male adult whale shark (total length, 8.8 m; Individual ID, sRhiTyp1) was sampled at Okinawa Churaumi Aquarium and used for genomic DNA extraction. Extraction of ultrahigh molecular weight DNA was performed by collecting blood cells by centrifugation, and the collected cells were embedded in agarose plugs (4.0 × 105 cells/plug). The plugs were prepared and processed with the CHEF Mammalian Genomic DNA Plug Kit (Bio-Rad 1703591). High DNA integrity was ensured with pulse-field gel electrophoresis. These animals were introduced into the aquarium in accordance with local regulations before those species were assessed as endangered. Animal handling and sample collections at the aquarium were conducted by veterinary staff without restraining the individuals under the experiment ID AT19002 approved by the Institutional Animal Care and Use Committee of the Okinawa Churashima Foundation in accordance with the Husbandry Guidelines approved by the Ethics and Welfare Committee of the Japanese Association of Zoos and Aquariums. All other experiments were conducted in accordance with the Guideline of the Institutional Animal Care and Use Committee (IACUC) of RIKEN Kobe Branch (Approval ID: H16-11).

### Genome sequencing and assembly

The DNA extracted as described above was subjected to library preparation using Ultra-Long DNA Sequencing Kit (Oxford Nanopore Technologies, SQK-ULK001). The prepared library was sequenced with R9.4.1 (FLO-PRO002) on PromethION. The raw sequencing output was processed with a basecaller Guppy Ver. 5 or 6 with Super-accurate basecalling mode.

Long reads were assembled using Flye v2.9, Racon v1.4.22, and Medaka v1.4.3 [41,42], resulting in 3,350 contigs with an N50 length of 17,247,868 bases. The contigs were then polished with the short-read data using FMLRC2 [43]. Hi-C reads from [7](SRX15207325 and SRX15207324) were mapped to the polished contigs with HiC-Pro v3.1.0 with the default parameter [44], after building an index with Bowtie2 v2.4.4 and SAMtools v1.12 [45,46]. Scaffolding was performed using YaHS v1.2 with the -r 2000,5000,10000,20000,50000,100000,200000,500000,1000000,2000000,5000000,100000 00,20000000,50000000,100000000,200000000,500000000 [47]. We manually curated the contact map using Juicebox v2.17.00 [48–50] (Figure 1B).

To ensure consistency in sequence orientation with the previous genome assembly version (sRhiTyp1.1), we aligned the direction of the sequences against the former version using minimap2 v2.24 with the -x asm5 and --secondary=no options [51]. Based on the resulting PAF file, the orientation of each scaffold was determined by identifying the alignment strand with the highest total alignment length to a single reference scaffold. When no unique best alignment was available, the orientation was maintained.

To compare the new and old assemblies, we aligned our new assembly to the former version (sRhiTyp1.1) as a reference using minimap2 with the -x asm5 option [51]. Based on the alignment results, dot plots were generated using a customized version of the dotPlotly script with the options -m 1000 -q 1000 -s -l -x (https://github.com/tpoorten/dotPlotly).

The assembly was evaluated using gVolante v2.0.0 and BUSCO v5.7.0 with the vertebrata_odb10 database [52,53].

### Gene prediction

We constructed a repeat model for each strain using RepeatModeler v2.0.5 with the - LTRStruct option and identified repeat sequences using RepeatMasker v4.1.5 with the options -nolow -xsmall [54,55]. Next, we first performed training for Augustus using BUSCO v5.7.0_cv1 with the -augustus and -long option based on the vertebrata_odb10 dataset [53]. We then predicted the protein-coding exons using Augustus v3.2.3 with the options --softmasking=1 --alternatives-from-evidence=true and our own config file, incorporating three hint files derived from transcript evidence, homolog protein evidence, and repetitive regions [56,57]. To generate the transcript-based hint file, we performed adapter trimming to raw reads from RNA-seq using fastp [58] and mapped the trimmed reads to the assembly using HISAT2 v2.2.1 [59] (accession numbers are shown in Table S2). For a protein hint file, we mapped peptide data from *Stegostoma tigrinum* (GCF_030684315.1, [60]) to the assembly using Exonerate v2.4.0 [61]. For a hint file from repeat regions, we used the RepeatMasker output. These outputs were converted to the appropriate format as instructed by the developer of Augustus (https://bioinf.uni-greifswald.de/bioinf/wiki/pmwiki.php?n=Augustus.Augustus).

### Identification of sex chromosomes

Short reads from female and male genomes were mapped to the scaffolds using BWA2 v2.2.1 with default parameters [62] (accession numbers are shown in Table S2). The read depth was calculated with SAMtools v1.19 and BEDtools v2.31.1 [46,63], then normalized by the mean depth across the entire genome. We compared the female and male read depths and classified scaffolds with a female-to-male depth ratio of approximately two as putative X chromosome (Figure 2A, C). One such chromosome, scaffold0040, was confirmed to be partially identical to a previously reported X chromosome [7] (Figure 2B). To identify potential Y scaffolds, we searched for scaffolds meeting the criteria of a female-to-male depth ratio < 0.1, a female depth < 0.l, a male depth > 0.3 and a length > 100 Kbp. Based on these thresholds, we identified two scaffolds, scaffold0112 (645 Kbp) and scaffold0679 (195 Kbp) (Figure 2D). We then examined the tandem repeats in these putative Y scaffolds using ModDotPlot v0.8.7 [64] (Figure 2E, F).

### Calculation of synonymous and non-synonymous substitution rate

We constructed ortholog groups (OGs) from the peptide sequences of whale shark(*R. typus*, this study), zebra shark (*Stegostoma tigrinum*, GCF_030684315.1, [60]), whitespotted bamboo shark (*Chiloscyllium plagiosum*, GCF_004010195.1, [65]), thorny skate (*Amblyraja radiata*, GCF_010909765.2, [66]) and elephant fish (*Callorhinchus milii*, GCF_018977255.1, [67]) using SonicParanoid v2.0.4 with default parameters [68], after selecting the longest isoform per gene using the agat_sp_keep_longest_isoform.pl script from AGAT v1.0.0 [69]. We then inferred phylogenetic trees for each OG using IQ-TREE v2.3.6 with the default parameters, after aligning the corresponding nucleotide sequences based on peptide alignments with MAFFT v7.526 and tranalign program of EMBOSS v6.6.0.0 [70–72]. Based on these phylogenetic trees, we used ETE3 v3.1.2 to identify OGs in which *R. typus* and *S. tigrinum* formed a monophyletic clade containing single-copy genes, resulting in 11,556 such OGs [73]. Next, for each OG, we calculated the *d*_*S*_ and *d*_*N*_ between these two species based on the YN model using the codeml program in PAML v4.9 with the following options: runmode = -2, model = 2, fix_kappa = 0, kappa = 2, fix_omega = 0 and omega = 1 [74]. We excluded the genes with T < 0.01 or T > 2. When we compared *d*_*S*_ (or *d*_*N*_) between genes on autosomes and the X chromosome, we only utilized genes that were located on autosomes in both species (or on the X chromosome in both species), excluding four genes that had translocated between autosomes and the X chromosome.

## Data Availability

The assembled sequence and gene model have been deposited in the figshare (https://figshare.com/projects/ykawaguchi-jinta/262804)

## Authors’ contributions

YWK (Conceptualization [supporting], Analysis [lead], Methodology [lead], Visualization [lead], Writing—original draft [lead], Writing—review & editing [lead]), RM (Methodology [lead]) and SK (Conceptualization [lead], Analysis [supporting], Methodology [lead], Visualization [supporting], Writing—original draft [supporting], Writing—review & editing [supporting]).

## Acknowledgments

We are grateful to Ryo Nozu, Kiyomi Murakumo, and Keiichi Sato at Okinawa Churaumi Aquarium for assistance in animal sampling, and to Kazuaki Yamaguchi and Osamu Nishimura for assistance in genome sequencing. We also thank GeneBay, Inc. for sequencing and assembly.

## References

1. Li H, Durbin R. Genome assembly in the telomere-to-telomere era. Nat Rev Genet. Springer Science and Business Media LLC; 25:658–702024;

2. Howe K, Chow W, Collins J, Pelan S, Pointon D-L, Sims Y, et al. Significantly improving the quality of genome assemblies through curation. Gigascience. Oxford University Press (OUP); 10:giaa1532021;

3. Read TD, Petit RA 3rd, Joseph SJ, Alam MT, Weil MR, Ahmad M, et al. Draft sequencing and assembly of the genome of the world’s largest fish, the whale shark: Rhincodon typus Smith 1828. BMC Genomics. BioMed Central; 18:5322017;

4. Hara Y, Yamaguchi K, Onimaru K, Kadota M, Koyanagi M, Keeley SD, et al. Shark genomes provide insights into elasmobranch evolution and the origin of vertebrates. Nat Ecol Evol. Springer Science and Business Media LLC; 2:1761–712018;

5. Tatsumi K, Nishimura O, Itomi K, Tanegashima C, Kuraku S. Optimization and cost-saving in tagmentation-based mate-pair library preparation and sequencing. Biotechniques. Informa UK Limited; 58:253–72015;

6. Weber JA, Park SG, Luria V, Jeon S, Kim H-M, Jeon Y, et al. The whale shark genome reveals how genomic and physiological properties scale with body size. Proc Natl Acad Sci U S A. 117:20662–712020;

7. Yamaguchi K, Uno Y, Kadota M, Nishimura O, Nozu R, Murakumo K, et al. Elasmobranch genome sequencing reveals evolutionary trends of vertebrate karyotype organization. Genome Res. 33:1527–402023;

8. Nishimura O, Rozewicki J, Yamaguchi K, Tatsumi K, Ohishi Y, Ohta T, et al. Squalomix: shark and ray genome analysis consortium and its data sharing platform. F1000Res. F1000 Research Ltd; 11:10772022;

9. Kadota M, Nishimura O, Miura H, Tanaka K, Hiratani I, Kuraku S. Multifaceted Hi-C benchmarking: what makes a difference in chromosome-scale genome scaffolding? Gigascience. Oxford University Press (OUP); 2020; doi: 10.1093/gigascience/giz158.

10. Matsumoto R, Matsumoto Y, Ueda K, Suzuki M, Asahina K, Sato K. Sexual maturation in a male whale shark (Rhincodon typus) based on observations made over 20 years of captivity. Fish Bull (Wash DC). NMFS Publications Office; 117:78–862019;

11. Uno Y, Nozu R, Kiyatake I, Higashiguchi N, Sodeyama S, Murakumo K, et al. Cell culture-based karyotyping of orectolobiform sharks for chromosome-scale genome analysis. Commun Biol. Springer Science and Business Media LLC; 3:6522020;

12. Wu J, Liu F, Jiao J, Luo H, Fan S, Liu J, et al. Comparative genomics illuminates karyotype and sex chromosome evolution of sharks. Cell Genom. Elsevier BV; 4:1006072024;

13. Niwa T, Uno Y, Ohishi Y, Kadota M, Aburatani N, Kiyatake I, et al. Sharks and rays have the oldest vertebrate sex chromosome with unique sex determination mechanisms. Proc Natl Acad Sci U S A. Proceedings of the National Academy of Sciences; 122:e25136761222025;

14. Hara Y, Kuraku S. Intragenomic mutational heterogeneity: structural and functional insights from gene evolution. Trends Genet. Elsevier; 2025; doi: 10.1016/j.tig.2025.03.007.

15. Stapley J, Feulner PGD, Johnston SE, Santure AW, Smadja CM. Variation in recombination frequency and distribution across eukaryotes: patterns and processes. Philos Trans R Soc Lond B Biol Sci. The Royal Society; 2017; doi: 10.1098/rstb.2016.0455.

16. Meisel RP, Connallon T. The faster-X effect: integrating theory and data. Trends Genet. Elsevier BV; 29:537–442013;

17. Miyata T, Hayashida H, Kuma K, Mitsuyasu K, Yasunaga T. Male-driven molecular evolution: a model and nucleotide sequence analysis. Cold Spring Harb Symp Quant Biol. Cold Spring Harbor Laboratory; 52:863–71987;

18. Ellegren H. Characteristics, causes and evolutionary consequences of male-biased mutation. Proc Biol Sci. The Royal Society; 274:1–102007;

19. Li WH, Yi S, Makova K. Male-driven evolution. Curr Opin Genet Dev. Elsevier BV; 12:650–62002;

20. Charlesworth B, Campos JL, Jackson BC. Faster-X evolution: Theory and evidence from Drosophila. Mol Ecol. Mol Ecol; 27:3753–712018;

21. Axelsson E, Webster MT, Smith NGC, Burt DW, Ellegren H. Comparison of the chicken and turkey genomes reveals a higher rate of nucleotide divergence on microchromosomes than macrochromosomes. Genome Res. Cold Spring Harbor Laboratory; 15:120–52005;

22. Kawakami T, Smeds L, Backström N, Husby A, Qvarnström A, Mugal CF, et al. A high-density linkage map enables a second-generation collared flycatcher genome assembly and reveals the patterns of avian recombination rate variation and chromosomal evolution. Mol Ecol. Wiley; 23:4035–582014;

23. Tigano A, Khan R, Omer AD, Weisz D, Dudchenko O, Multani AS, et al. Chromosome size affects sequence divergence between species through the interplay of recombination and selection. Evolution. Wiley; 76:782–982022;

24. Jones GH, Franklin FCH. Meiotic crossing-over: obligation and interference. Cell. Elsevier BV; p. 246–8.

25. Lau Q, Igawa T, Ogino H, Katsura Y, Ikemura T, Satta Y. Heterogeneity of synonymous substitution rates in the Xenopus frog genome. PLoS One. Public Library of Science (PLoS); 15:e02365152020;

26. Sendell-Price AT, Tulenko FJ, Pettersson M, Kang D, Montandon M, Winkler S, et al. Low mutation rate in epaulette sharks is consistent with a slow rate of evolution in sharks. Nat Commun. Springer Science and Business Media LLC; 14:66282023;

27. Long DJ. Sharks from the la meseta formation (Eocene), Seymour island, antarctic peninsula. J Vertebr Paleontol. Informa UK Limited; 12:11–321992;

28. Church DM, Goodstadt L, Hillier LW, Zody MC, Goldstein S, She X, et al. Lineagespecific biology revealed by a finished genome assembly of the mouse. PLoS Biol. Public Library of Science (PLoS); 7:e10001122009;

29. Alföldi J, Di Palma F, Grabherr M, Williams C, Kong L, Mauceli E, et al. The genome of the green anole lizard and a comparative analysis with birds and mammals. Nature. Springer Science and Business Media LLC; 477:587–912011;

30. Donoghue PCJ, Benton MJ. Rocks and clocks: calibrating the Tree of Life using fossils and molecules. Trends Ecol Evol. Elsevier BV; 22:424–312007;

31. Montoya-Burgos JI. Patterns of positive selection and neutral evolution in the proteincoding genes of Tetraodon and Takifugu. PLoS One. Public Library of Science (PLoS); 6:e248002011;

32. Kuraku S, Feiner N, Keeley SD, Hara Y. Incorporating tree-thinking and evolutionary time scale into developmental biology. Development, Growth & Differentiation. John Wiley & Sons, Ltd; 58:131–422016;

33. Kong A, Gudbjartsson DF, Sainz J, Jonsdottir GM, Gudjonsson SA, Richardsson B, et al. A high-resolution recombination map of the human genome. Nat Genet. Springer Science and Business Media LLC; 31:241–72002;

34. Haenel Q, Laurentino TG, Roesti M, Berner D. Meta-analysis of chromosome-scale crossover rate variation in eukaryotes and its significance to evolutionary genomics. Mol Ecol. sJohn Wiley & Sons, Ltd; 27:2477–972018;

35. Supek F, Lehner B. Differential DNA mismatch repair underlies mutation rate variation across the human genome. Nature. Springer Science and Business Media LLC; 521:81–42015;

36. Schuster-Böckler B, Lehner B. Chromatin organization is a major influence on regional mutation rates in human cancer cells. Nature. Springer Science and Business Media LLC; 488:504–72012;

37. Charlesworth B, Coyne JA, Barton NH. The relative rates of evolution of sex chromosomes and autosomes. Am Nat. 130:113–461987;

38. Mank JE, Vicoso B, Berlin S, Charlesworth B. Effective population size and the Faster-X effect: empirical results and their interpretation. Evolution. Wiley; 64:663–742010;

39. Julien P, Brawand D, Soumillon M, Necsulea A, Liechti A, Schütz F, et al. Mechanisms and evolutionary patterns of mammalian and avian dosage compensation. PLoS Biol. Public Library of Science (PLoS); 10:e10013282012;

40. Mank JE, Nam K, Ellegren H. Faster-Z evolution is predominantly due to genetic drift. Mol Biol Evol. Oxford University Press (OUP); 27:661–702010;

41. Vaser R, Sović I, Nagarajan N, Šikić M. Fast and accurate de novo genome assembly from long uncorrected reads. Genome Res. 27:737–462017;

42. Kolmogorov M, Yuan J, Lin Y, Pevzner PA. Assembly of long, error-prone reads using repeat graphs. Nat Biotechnol. Springer Science and Business Media LLC; 37:540–62019;

43. Mak QXC, Wick RR, Holt JM, Wang JR. Polishing DE Novo nanopore assemblies of bacteria and eukaryotes with FMLRC2. Mol Biol Evol. Oxford University Press (OUP); 40:msad0482023;

44. Servant N, Varoquaux N, Lajoie BR, Viara E, Chen C-J, Vert J-P, et al. HiC-Pro: an optimized and flexible pipeline for Hi-C data processing. Genome Biol. Springer Science and Business Media LLC; 16:2592015;

45. Langmead B, Salzberg SL. Fast gapped-read alignment with Bowtie 2. Nat Methods. 9:357–92012;

46. Li H, Handsaker B, Wysoker A, Fennell T, Ruan J, Homer N, et al. The Sequence Alignment/Map format and SAMtools. Bioinformatics. academic.oup.com; 25:2078–92009;

47. Zhou C, McCarthy SA, Durbin R. YaHS: yet another Hi-C scaffolding tool. Bioinformatics. Oxford University Press (OUP); 39:btac8082023;

48. Rao SSP, Huntley MH, Durand NC, Stamenova EK, Bochkov ID, Robinson JT, et al. A 3D map of the human genome at kilobase resolution reveals principles of chromatin looping. Cell. Elsevier BV; 159:1665–802014;

49. Durand NC, Robinson JT, Shamim MS, Machol I, Mesirov JP, Lander ES, et al. Juicebox provides a visualization system for Hi-C contact maps with unlimited zoom. Cell Syst. 3:99–1012016;

50. Dudchenko O, Shamim MS, Batra SS, Durand NC, Musial NT, Mostofa R, et al. The Juicebox Assembly Tools module facilitates de novo assembly of mammalian genomes with chromosome-length scaffolds for under $1000. bioRxiv. bioRxiv;

51. Li H. Minimap2: pairwise alignment for nucleotide sequences. Bioinformatics. academic.oup.com; 34:3094–1002017;

52. Nishimura O, Hara Y, Kuraku S. gVolante for standardizing completeness assessment of genome and transcriptome assemblies. Bioinformatics. Oxford Academic; 33:3635–72017;

53. Manni M, Berkeley MR, Seppey M, Simão FA, Zdobnov EM. BUSCO update: Novel and streamlined workflows along with broader and deeper phylogenetic coverage for scoring of eukaryotic, prokaryotic, and viral genomes. Mol Biol Evol. Oxford University Press (OUP); 38:4647–542021;

54. Smit AFA, Hubley R. RepeatModeler Open-1.0. 2008-2015. Seattle, USA: Institute for Systems Biology. Available.

55. Smit AFA, Hubley R, Green P. RepeatMasker Open-4.0. 2013–2015.

56. Keller O, Kollmar M, Stanke M, Waack S. A novel hybrid gene prediction method employing protein multiple sequence alignments. Bioinformatics. 27:757–632011;

57. Stanke M, Diekhans M, Baertsch R, Haussler D. Using native and syntenically mapped cDNA alignments to improve de novo gene finding. Bioinformatics. Oxford University Press (OUP); 24:637–442008;

58. Chen S, Zhou Y, Chen Y, Gu J. fastp: an ultra-fast all-in-one FASTQ preprocessor. Bioinformatics. 34:i884–902018;

59. Kim D, Langmead B, Salzberg SL. HISAT: a fast spliced aligner with low memory requirements. Nat Methods. nature.com; 12:357–602015;

60. Lee S-H, Fedrigo O, Soler-Clavel L, Humble E, Lesturgie P, Balacco J, et al. Insights into the evolution of ancient shark and ray sex chromosomes. bioRxiv.

61. Slater GSC, Birney E. Automated generation of heuristics for biological sequence comparison. BMC Bioinformatics. 6:312005;

62. Vasimuddin M, Misra S, Li H, Aluru S. Efficient architecture-aware acceleration of BWA-MEM for multicore systems. 2019 IEEE International Parallel and Distributed Processing Symposium (IPDPS). IEEE; p. 314–24.

63. Quinlan AR, Hall IM. BEDTools: a flexible suite of utilities for comparing genomic features. Bioinformatics. 26:841–22010;

64. Sweeten AP, Schatz MC, Phillippy AM. ModDotPlot-rapid and interactive visualization of tandem repeats. Bioinformatics. Oxford University Press (OUP); 40:btae4932024;

65. Zhang Y, Gao H, Li H, Guo J, Ouyang B, Wang M, et al. The white-spotted bamboo shark genome reveals chromosome rearrangements and fast-evolving immune genes of cartilaginous fish. iScience. Elsevier BV; 23:1017542020;

66. Rhie A, McCarthy SA, Fedrigo O, Damas J, Formenti G, Koren S, et al. Towards complete and error-free genome assemblies of all vertebrate species. Nature. Springer Science and Business Media LLC; 592:737–462021;

67. Nakatani Y, Shingate P, Ravi V, Pillai NE, Prasad A, McLysaght A, et al. Reconstruction of proto-vertebrate, proto-cyclostome and proto-gnathostome genomes provides new insights into early vertebrate evolution. Nat Commun. 12:44892021;

68. Cosentino S, Sriswasdi S, Iwasaki W. SonicParanoid2: fast, accurate, and comprehensive orthology inference with machine learning and language models. Genome Biol. Springer Science and Business Media LLC; 25:1952024;

69. Dainat J. Another Gtf/Gff Analysis Toolkit (AGAT): Resolve interoperability issues and accomplish more with your annotations.

70. Minh BQ, Schmidt HA, Chernomor O, Schrempf D, Woodhams MD, von Haeseler A, et al. IQ-TREE 2: New models and efficient methods for phylogenetic inference in the genomic era. Mol Biol Evol. Oxford University Press (OUP); 37:1530–42020;

71. Katoh K, Standley DM. MAFFT multiple sequence alignment software version 7: improvements in performance and usability. Mol Biol Evol. 30:772–802013;

72. Rice P, Longden I, Bleasby A. EMBOSS: The European molecular biology open software suite. Trends Genet. Elsevier BV; 16:276–72000;

73. Huerta-Cepas J, Serra F, Bork P. ETE 3: Reconstruction, Analysis, and Visualization of Phylogenomic Data. Mol Biol Evol. 33:1635–82016;

74. Yang Z. PAML 4: phylogenetic analysis by maximum likelihood. Mol Biol Evol. Oxford University Press (OUP); 24:1586–912007;

